# Generation of a Panel of Induced Pluripotent Stem Cells From Chimpanzees: a Resource for Comparative Functional Genomics

**DOI:** 10.1101/008862

**Authors:** Irene Gallego Romero, Bryan J. Pavlovic, Irene Hernando-Herraez, Nicholas E. Banovich, Courtney L. Kagan, Jonathan E. Burnett, Constance H. Huang, Amy Mitrano, Claudia I. Chavarria, Inbar Friedrich Ben-Nun, Yingchun Li, Karen Sabatini, Trevor Leonardo, Mana Parast, Tomas Marques-Bonet, Louise C. Laurent, Jeanne F. Loring, Yoav Gilad

## Abstract

Comparative genomics studies in primates are extremely restricted because we only have access to a few types of cell lines from non-human apes and to a limited collection of frozen tissues. In order to gain better insight into regulatory processes that underlie variation in complex phenotypes, we must have access to faithful model systems for a wide range of tissues and cell types. To facilitate this, we have generated a panel of 7 fully characterized chimpanzee (*Pan troglodytes*) induced pluripotent stem cell (iPSC) lines derived from fibroblasts of healthy donors. All lines appear to be free of integration from exogenous reprogramming vectors, can be maintained using standard iPSC culture techniques, and have proliferative and differentiation potential similar to human and mouse lines. To begin demonstrating the utility of comparative iPSC panels, we collected RNA sequencing data and methylation profiles from the chimpanzee iPSCs and their corresponding fibroblast precursors, as well as from 7 human iPSCs and their precursors, which were of multiple cell type and population origins. Overall, we observed much less regulatory variation within species in the iPSCs than in the somatic precursors, indicating that the reprogramming process has erased many of the differences observed between somatic cells of different origins. We identified 4,918 differentially expressed genes and 3,598 differentially methylated regions between iPSCs of the two species, many of which are novel inter-species differences that were not observed between the somatic cells of the two species. Our panel will help realise the potential of iPSCs in primate studies, and in combination with genomic technologies, transform studies of comparative evolution.

## Introduction

Comparative functional genomic studies of humans and other primates have been consistently hindered by a lack of samples (Gallego Romero et al. 2012). In spite of their clear potential to inform our understanding of both human evolution and disease, practical and ethical concerns surrounding working with non-human primates have constrained the field to using a limited set of cell types collected non-invasively, primarily lymphoblastoid cell lines (LCLs) and fibroblasts. Comparative studies of any other primate tissue have been limited to using post-mortem (typically frozen) materials, thereby precluding most experimental manipulation and yielding primarily observational insights (see, for example, Blekhman et al. 2010; Blekhman et al. 2008; Brawand et al. 2011).

An alternative has been to use model organisms in an attempt to recapitulate inter-primate regulatory differences. The typical approach involves the introduction of sequences of evolutionary interest into a model system, and then searching for spatial or temporal differences in gene expression that can be ascribed to the introduced sequence (Cotney et al. 2013; Enard et al. 2009). This is a difficult and challenging approach and, perhaps as a result, there are still only a handful of well-described examples of human-specific regulatory adaptations in primates (McLean et al. 2011; Prabhakar et al. 2008) and even fewer cases where the underlying regulatory mechanisms have been resolved (Pollard et al. 2006; Rockman et al. 2005). While these studies are useful and often informative, they also entail assumptions of functional conservation across the model system and the species of interest that may not necessarily be true (Gallego Romero et al. 2012).

Induced pluripotent stem cells (iPSCs) can provide a viable means of circumventing these concerns and limitations, at least with respect to a subset of phenotypes (those that can be studied in *in-vitro* systems). Reprogramming somatic cell lines to a stable and self-sustaining pluripotent state (Takahashi et al. 2007; Takahashi and Yamanaka 2006) has become routine practice for human and murine cell lines, but extension to other animals, especially non-human primates, is not yet widespread despite some notable exceptions (Ben-Nun et al. 2011; Marchetto et al. 2013b). Instead, the broadest application of iPSCs to date has been the generation of lines derived from patients suffering from any of a sizable number of genetic disorders (Cohen and Melton 2011; Israel et al. 2012; Liu et al. 2012; Merkle and Eggan 2013; Wang et al. 2014) with the twin aims of providing a deeper understanding of disease phenotypes as well as highlighting new therapeutic avenues. These lines have been shown to mimic some relevant patient phenotypes observed *in vivo*, both as iPSCs and when differentiated into other pertinent cell types, and thus suggest a way forward for regenerative medical applications; but more generally, they also highlight the tantalizing flexibility of iPSCs as a means of exploring many different developmental and cell lineage determination pathways.

Thus, the development of an iPSC system for comparative genomic studies in primates will allow us to compare regulatory pathways and complex phenotypes in humans and our close evolutionary relatives using appropriate models for different tissues and cell types. This will be a powerful resource with which to examine the contribution of changes in gene regulation to human evolution and diversity. To demonstrate the validity of this approach, we have generated a panel of 7 chimpanzee iPSC lines that are fully characterized and comparable to human iPSC lines in their growth and differentiation capabilities.

## Results

We generated a panel of induced pluripotent stem cell lines from seven chimpanzees through electroporation of episomal plasmids expressing *OCT3/4* (also known as *POU5F1*), *SOX2, KLF4, L-MYC, LIN28*, and an shRNA *p53* knockdown (Okita et al. 2011), as well as an *in vitro*-transcribed *EBNA1* mRNA transcript (Chen et al. 2011; Howden et al. 2006) that promotes increased exogenous vector retention in the days following electroporation. Our chimpanzee panel is comprised of seven healthy individuals (4 female, 3 male, further details on these individuals are given in supplementary table 1) ranging from 9 to 17 years old. Fibroblasts from 5 of the 7 individuals were purchased from the Coriell Cell Repositories while the remaining two (C40210, C40280) were derived from 3 mm skin punch biopsies directly collected from animals at the Yerkes Primate Research Center of Emory University.

### Characterizing the chimpanzee iPSCs

The chimpanzee iPSC lines closely resembled human iPSC lines in morphology (figure 1a; all images shown in main text are from chimpanzee line C4955. Similar images of the other lines are available as supplementary figures 1 to 7). The lines could be maintained in culture for at least 60 passages without loss of pluripotency or self-renewal capability using standard iPSC culture conditions, both on mouse embryonic fibroblast (MEF) feeder cells and in feeder-free conditions. The genomes of all our lines appear stable; all exhibited normal karyotypes after reprogramming and more than 15 passages in culture, ruling out the presence of gross chromosomal abnormalities that are sometimes reported to be introduced during the reprogramming and culture process (figure 1b, supplementary figure 1).

**Figure 1:**
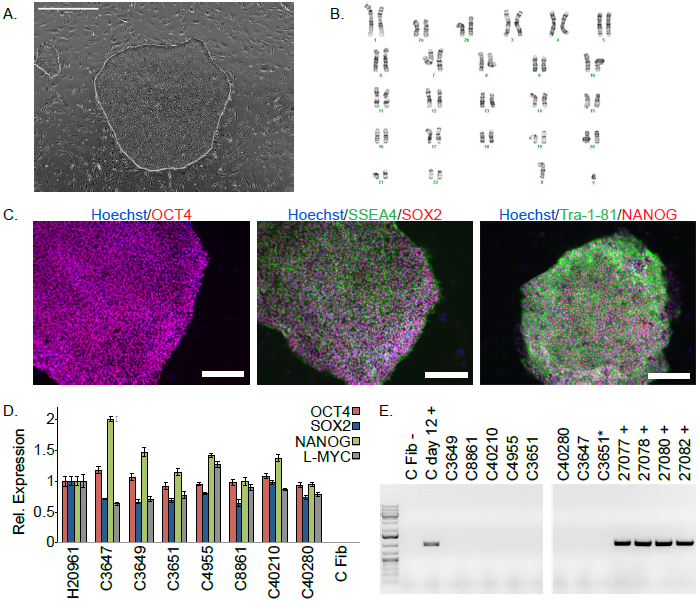
Characterization of chimpanzee iPSC lines. a. Phase contrast image of representative chimpanzee iPSC line. Scale bar: 1000 μm. b. Representative karyotype from chimpanzee iPSC line after > 15 passages, showing no abnormalities. c. ICC staining of iPSC lines with antibodies for pluripotency markers as indicated. Scale bar: 200 μm. d. Quantitative PCR testing for expression of endogenous pluripotency factors in all 7 chimpanzee iPSC lines. Line H20961 is a male human iPSC line generated in-house used as reference. e. PCR gel showing an absence of exogenous episomal reprogramming factors in all 7 chimpanzee iPSC lines. All PCRs were carried out on templates extracted from passage >15 with the exception of C3651*, which is from passage 2. Fib – is a negative fibroblast control (from individual C8861) prior to transfection, day 12 + is a positive control 12 days after transfection, 27077 + to 27082 + are the plasmids used in transfection.

We confirmed nuclear expression of *OCT3/4, SOX2* and *NANOG* in all lines by immunocytochemistry (figure 1c; supplementary figure 2). The pluripotent cells also expressed the surface antigens Tra-1-81 and SSEA4, while cells collected from the center of differentiating colonies expressed SSEA1 at levels comparable to differentiating colonies of human iPSC lines (supplementary figure 3). To confirm that the observed pluripotency expressed genes were of endogenous origin, we performed qPCR with primers designed to specifically amplify the endogenous *OCT3/4, SOX2, NANOG* and *L-MYC* transcripts (figure 1d; all PCR primers used in this work are listed in supplementary table 2). Indeed, we found no evidence of exogenous gene expression after 10 passages (supplementary figure 4), and no traces of genomic integration or residual episomal plasmid retention after 15 passages (figure 1e). These observations strongly indicate that self-renewal in our chimpanzee iPSC lines is maintained solely through endogenous gene expression.

To confirm pluripotency and test the differentiation capabilities of our lines we generated embryoid bodies and assayed their ability to spontaneously differentiate into the three main germ layers by immunocytochemistry (figure 2a; supplementary figure 5). Additionally, we carried out teratoma formation assays in four of the lines using Fox Chase SCID-beige and CB17.Cg-*Prkdc*^*scid*^*Lyst*^*bg-J*^/Crl immunodeficient male mice. All four iPSC lines were capable of generating tumours in mice, and all tumours examined contained tissues of endodermal, ectodermal and mesodermal origins (figure 2b, supplementary figure 6). To confirm the chimpanzee origin of these tissues, we extracted and Sanger-sequenced mitochondrial DNA from the tumour (supplementary figure 7). Finally we further characterized pluripotency in our lines by the application of PluriTest, a bioinformatic classifier that compares the array-based estimates of gene expression profiles of new lines to those obtained from over 400 well-characterized human pluripotent and terminally differentiated lines (Müller et al. 2011), modified to accommodate data from both species. All chimpanzee lines have pluripotency scores greater than the pluripotency threshold value of 20 (figure 3a, supplementary table 1).

**Figure 2:**
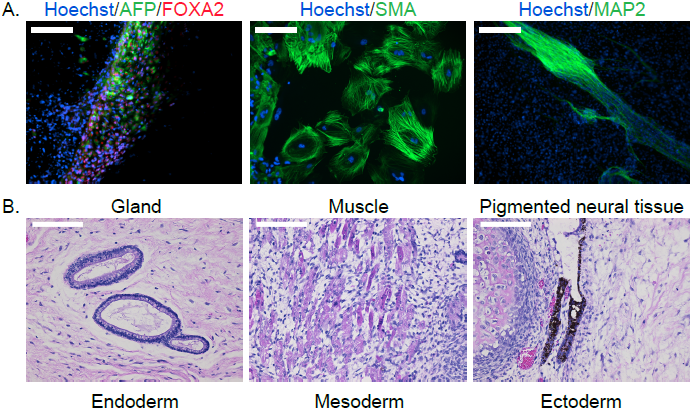
a. ICC staining of differentiated embryoid bodies with antibodies for the three germ layers as indicated. Scale bar: 200 μm. b. Histological staining of teratomas derived from iPSC line C4955, showing generation of tissues from all three germ layers. Scale bar: 100 μm.

**Figure 3:**
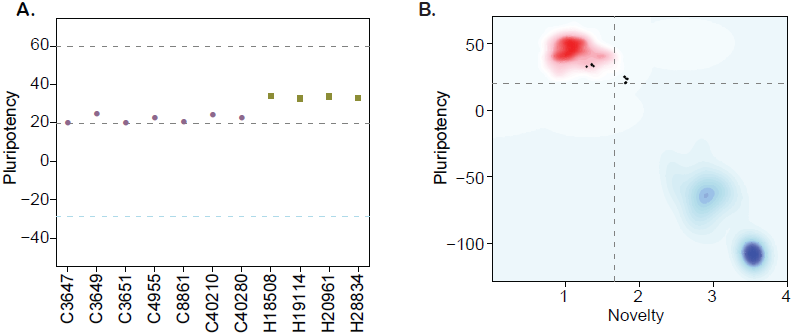
a. PluriTest pluripotency scores in the 7 chimpanzee lines and 4 human reference iPSC lines. Purple circles denote chimpanzees; yellow squares, humans. b. PluriTest results after removal of probes not mapping to the chimpanzee genome. All samples in the top left quadrant are human and have satisfactory pluripotency and novelty scores. Samples in the top right quadrant correspond to our chimpanzee iPSC panel, and have consistently high pluripotency yet high novelty scores.

We also calculated PluriTest novelty scores for all our samples. In human iPSCs, novelty values above 1.67 are suggestive of chromosomal duplications or expression of differentiation-associated genes. Human iPSCs with high novelty scores are typically either difficult to maintain and expand in culture (because they differentiate spontaneously at a high rate), or are not capable to differentiate well to all three germ layers. All of our chimpanzee lines had novelty scores above the 1.67 threshold (figure 3b). Yet, our chimpanzee lines can easily be maintained in culture on the one hand, and can be differentiated into all three germ layer lineages (in embryoid bodies in vitro and in teratomas in vivo) on the other hand. We thus hypothesize that the observed high novelty scores are likely driven by inter-species gene regulatory differences that the PluriTest assay, which was trained exclusively on human samples, interpreted as abnormal gene expression.

### Comparative analysis of gene expression and methylation data from iPSCs

To better examine gene expression and regulatory differences between human and chimpanzee iPSCs, we generated genome-wide RNA-sequencing and DNA methylation data (see methods) from all chimpanzee iPSC lines, as well as from 7 human iPSC lines also generated and validated in our laboratory. While all of our chimpanzee donors were from one subspecies and were all derived from fibroblasts (supplementary table 1), the human samples include both Caucasian and Yoruba individuals, and the iPSCs were generated from both fibroblasts and immortalised lymphoblastoid cell lines (LCLs; see supplementary table 3 for additional details). We designed the comparative study this way in order to demonstrate that regulatory differences between human and chimpanzee iPSCs cannot be explained by technical differences due to culturing conditions or the origin of the precursor cells.

To prevent biases due to genetic divergence between the two species, we chose to restrict our gene expression analyses to a curated set of genes with one-to-one orthology between humans and chimpanzees (Blekhman 2012; Blekhman et al. 2010). Following assessment of quality control metrics (see methods), we obtained normalised RPKM estimates for 12,503 genes that were expressed in at least 4 iPSC lines from either one of the species (see methods). We similarly restricted our DNA methylation analyses to a set of 335,307 high quality probes with a high degree of sequence conservation between humans and chimpanzees (as in Hernando-Herraez et al. 2013; see methods).

To examine broad patterns in the data, we used principal component analysis (PCA). We observed clear and robust separation of human and chimpanzee iPSC lines along the first principal component (PC), regardless of the data type used (figures 4a, 4b; Regression of PC1 by species; *P* < 10^−14^ for the expression data; *P* < 10^−12^ for the DNA methylation data). Within the human samples, PC2 appears to be driven by ethnicity, as we observe all Caucasian samples consistently clustering together despite their multiple cell types of origin (*P* = 0.005 for the association between PC2 and human ethnicity in the expression data, *P* = 0.046 in the methylation data).

**Figure 4:**
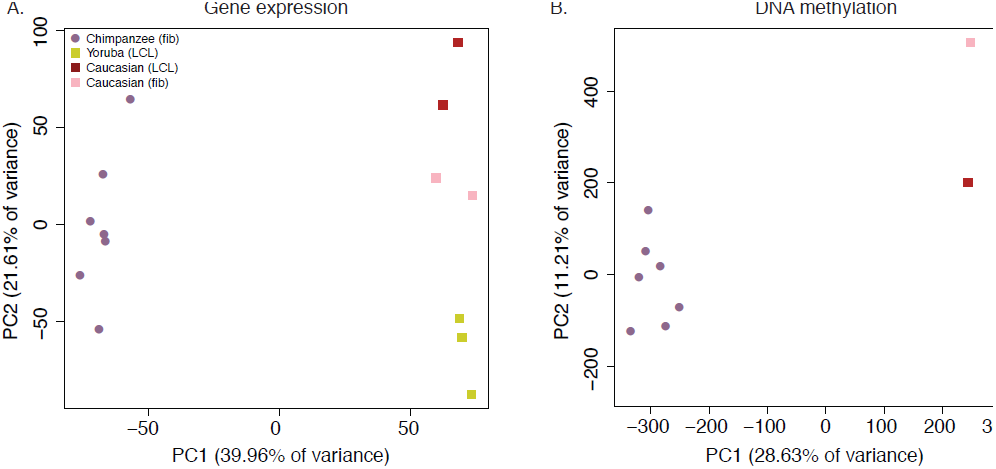
Principal component analysis plots of data from the iPSCs. a. PCA generated from expression data of 13,756 orthologous genes. b. PCA generated from DNA methylation data measured by 335,003 filtered probes.

We then analysed regulatory differences between the species by first focusing on the gene expression data. At an FDR of 1%, we identified 4,918 genes (39.3%) as differentially expressed (DE) between the iPSCs of the two species (supplementary table 4), a large number of which exhibit considerable inter-species fold changes in expression levels (supplementary figure 8). An analysis of functional annotation of the DE genes is not very informative, although 10 Gene Ontology (GO) Biological Process terms (Ashburner et al. 2000) are significantly overrepresented among these genes at an FDR of 5% (supplementary table 5). Enrichment in most of these terms is primarily driven by decreased expression of a group of ribosomal proteins in chimpanzee iPSCs relative to their human orthologs (supplementary table 4, supplementary figure 9). Additionally, we tested for concordance between our list of DE genes and a list of 2,730 genes that were previously classified as differentially expressed between human and non-human primate iPSC lines (Marchetto et al. 2013b). Given our stringent approach to consider orthologous genes, only 2,093 (77% of the 2,730) genes could be potentially analysed across the two studies. Of these, 1,481 genes are detectably expressed in our lines, and 1,079 (72.9%) are classified as DE between the species in both data sets (a highly significant enrichment; χ^2^ *P* < 10^−16^). Expression trends within these DE genes are in the same direction in both data sets in 1,062 of cases (98.42%).

Next, we used a similar approach to identify differentially methylated probes and regions between the iPSCs of both species (see methods). We identified 64,047 probes that are differentially methylated (DM) between the two species (at an FDR of 1%), 26,718 of which have a mean intergroup β difference ≥ 0.1, our arbitrary effect size threshold for retaining probes for differentially methylated region (DMR) identification and downstream analyses. Of these, 10,855 probes could be further grouped into 3,598 regions of 2 or more DM probes within 1 kb, which we designated DMRs; supplementary table 6); the numbers of probes and regions identified as DM at a range of mean interspecies β thresholds are given in supplementary table 7.

In order to consider the methylation and gene expression data jointly, we focused on a subset of 2,401 DMRs that could be associated with a single ENSEMBL gene (see methods). Overall, these DMRs were associated with 2,194 genes, of which 1,385 were also detectably expressed in the iPSCs, and 576 (41.6%) were classified as differentially expressed between the species, a slightly higher proportion than expected by chance alone (*P* = 0.1). We further classified the DMRs as either ‘promoter’, ‘genic’ or ‘mixed’ depending on their position relative to annotated gene transcripts (see methods). The overall set of DMRS, and DMRs found in gene bodies are significantly associated with 20 and 92 GO BP terms respectively after FDR correction, including terms related to neurogenesis and skeletal system development. Enrichment of several terms related to neurogenesis and skeletal system development is likewise marginally significant amongst promoter and mixed DMRs (supplementary table 8).

### Gene expression and methylation in iPSCs and their precursors

We additionally collected RNA-sequencing and methylation data from all cell lines used to generate both the chimpanzee and human iPSCs (supplementary table 9). Following quality control and normalisation steps we obtained RPKM values for 13,486 genes across all 28 iPSC and precursor samples (see methods). We also obtained DNA methylation profiles using the same 335,307 array probes we considered above from all samples. We performed PCA and found that the first PC was now significantly associated with tissue type in both data sets (*P* < 10^−29^ for the expression data; *P* < 10^−17^ for the methylation data; see figure 5 and supplementary tables 10 and 11). Data from the iPSCs of both species are clearly separated from data from the precursor somatic cells. Humans and chimpanzee samples are separated along PC2 (*P* < 10^−3^ for the expression data; *P* < 10^−4^ for the methylation data).

**Figure 5:**
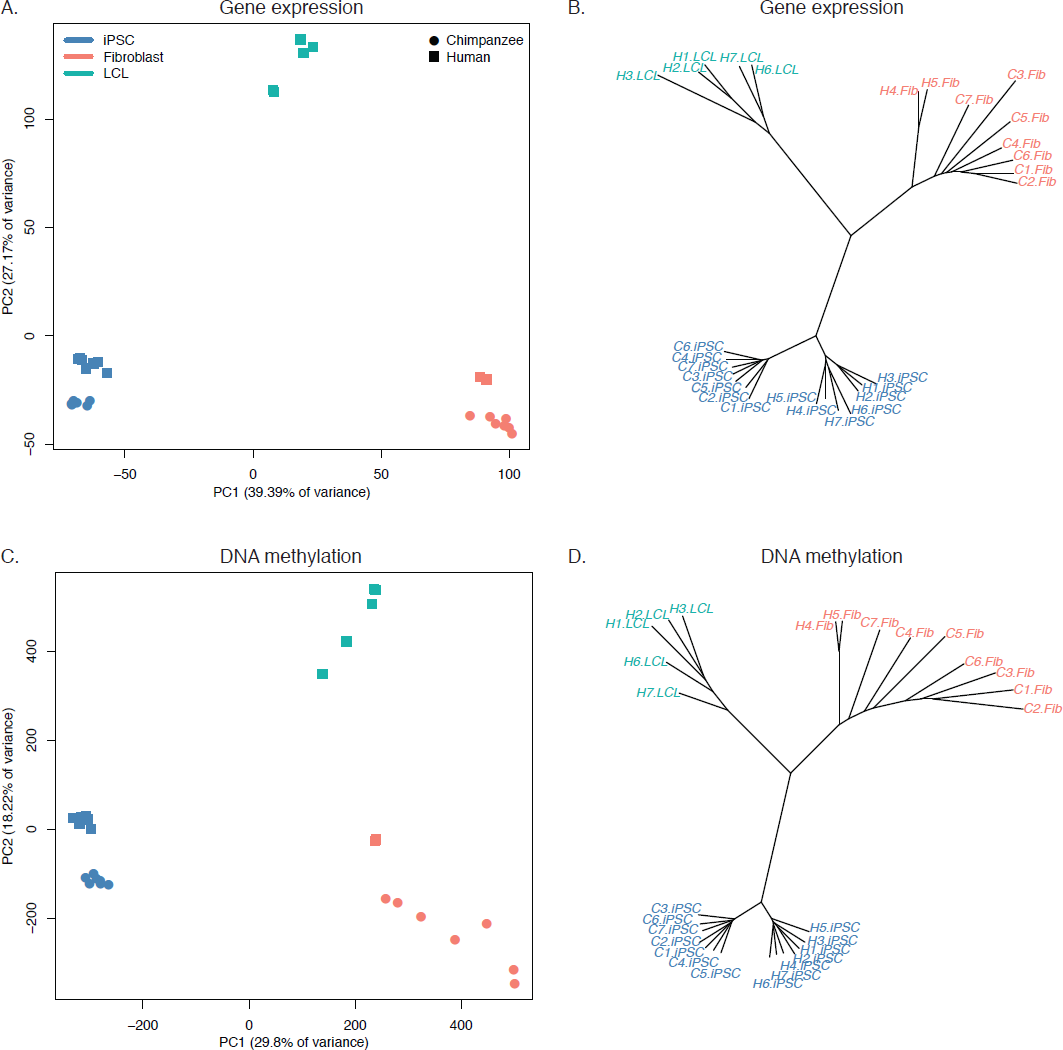
Relationships of iPSCs to their precursors. a. PCA of gene expression data from all iPSCs and their precursor cell lines. b. Neighbour-joining tree of Euclidean distances between all samples generated based on the gene expression data. c. PCA of DNA methylation data from all iPSCs and their precursor cell lines. d. Neighbour-joining tree of Euclidean distances between all samples generated based on the DNA methylation data.

Chimpanzee iPSCs have significantly overall higher levels of DNA methylation compared to their precursor somatic cells (*P* < 10^−15^; supplementary figure 10), an observation that extends to all genomic features we tested (supplementary figure 11). This is not surprising as a similar pattern has been previously shown in human pluripotent cell lines (Bock et al. 2011; Nazor et al. 2012). Remarkably, both DNA methylation and gene expression levels in iPSCs are relatively homogeneous within species, far more so than in their corresponding precursor cells (figures 5b, 5d: *P* < 10^−14^ when comparing overall pairwise distances within all chimpanzee iPSCs and within all chimpanzee fibroblasts in the methylation data; *P* < 10^−9^ for the same comparison in the gene expression data). Methylation levels in iPSCs also have significantly reduced coefficients of variation relative to their precursor lines (range of CVs for chimpanzee iPSCs = 0.78 - 0.80, for chimpanzee fibroblasts = 0.87 - 0.90; *P* < 10^−06^). We observed the same pattern in the human data, although in this case the multiple origins of the precursor cells contribute to the higher variation.

We then performed analyses of gene expression and methylation differences in the combined iPSC and precursor cells. First, we considered a comparison of the iPSCs and the precursor cells within species (see methods) and classified 9,495 genes as DE between chimpanzee fibroblasts and the corresponding iPSCs, whereas in humans the number of DE genes is 8,077 if we consider all iPSC lines and their precursors, 8,427 if we only consider those derived from LCLs (n = 5), and 5,617 if we only consider those derived from fibroblasts (n = 2; supplementary table 12). Similarly, we identified 18,029 DMRs between chimpanzee fibroblasts and iPSCs, and 12,078 DMRs between all human precursors and all human iPSCs (supplementary tables 13-14).

Next, we focused on a comparison of inter-species differences in gene expression and methylation levels across cell types. Following normalization and modelling of data from all samples (see methods), we classified 6,062 genes as DE between the chimpanzee precursor fibroblasts and the collection of human precursor LCLs and fibroblasts, as well as 84,747 DM probes and 9,107 DMRs (always at an FDR of 1%). Most of these regulatory differences, however, reflect variation across cell types rather than across species (6,437 genes and 70,312 probes are DE or DM between the human fibroblasts and LCLs, respectively). We thus considered only data from the fibroblast precursors in the two species. Although the sample size is considerably smaller (only 2 of the human iPSCs were reprogrammed from fibroblasts), we were able to identify 1,612 DE genes and 25,456 DM probes between human and chimpanzee fibroblasts, and 1,514 DE genes and 16,392 DM probes between the corresponding iPSCs of the two species.

Interestingly, although the overlap of inter-species DE genes and DM probes between the iPSCs and the precursors is considerable (37.7% of DE genes and 27.0% of DM probes), a large number of regulatory differences are only observed between the iPSC lines of the two species (supplementary figure 12). This observation is robust with respect to different approaches to normalize and model the data (supplementary figure 13), strongly suggesting that many of the differences we observe between our chimpanzee and human iPSC lines may be intrinsic features of the pluripotent state in these two species.

## Discussion

Induced pluripotent stem cells have the potential to transform our understanding of the biology of non-model organisms and facilitate functional comparative studies. To this end, we have generated a panel of 7 fully characterized chimpanzee iPSCs. All lines are capable of spontaneously giving rise to the three tissue germ layers *in vitro* and meet all currently established thresholds for pluripotency. The chimpanzee iPSC lines provide a tantalising avenue for investigating how changes in gene expression and regulation underlie the architecture of complex phenotypic traits in humans and our closest living relatives (Gallego Romero et al. 2012; Marchetto et al. 2013a). In particular, we believe that through the use of directed differentiation protocols, functional studies could be performed in cell types where strong *a priori* hypotheses support a role for selective pressure underlying inter-species divergence (e.g. liver, heart, kidney (Blekhman et al. 2010; Blekhman et al. 2008)). In that sense, we hope that this panel will be a useful tool to researchers interested in overcoming current limitations of comparative studies in primates. Crucially for this, all lines from the panel are readily available for sharing with investigators who may wish to address questions beyond the scope of our own research.

Other groups have previously generated pluripotent stem cells from primates (Ben-Nun et al. 2011; Chan et al. 2010; Deleidi et al. 2011; Hong et al. 2014; Liu et al. 2008; Okamoto and Takahashi 2011; Tomioka et al. 2010; Wu et al. 2012; Wu et al. 2010; Wunderlich et al. 2014). Indeed, a recent publication (Marchetto et al. 2013b) reported the generation of two chimpanzee and two bonobo (*Pan paniscus*) iPSC lines through the use of retroviral vectors. However, in the course of our work we have found that retroviral vector silencing in chimpanzee iPSCs was not as stable as in human iPSC lines generated at the same time through the same method (see Methods). Our use of episomal vectors circumvents this problem, and more broadly the problems of both random exogenous gene reactivation and disruption of the host genome through retroviral integration (Sommer et al. 2012).

More generally, while the sum total of primate PSC generation efforts so far has resulted in a sizable number of lines being established from various donors and species, it has also entailed the use of various reprogramming protocols and source cell types. We have generated iPSCs from a panel of seven individuals using a consistent protocol and cell type of origin. Given the panel size, it is powerful enough to robustly detect inter-species differences in gene expression, splicing and regulation. The fact that our panel contains both female and male lines also allows for future studies of sex-specific differences in gene expression in various cell types. Indeed, we have previously shown that this can be accomplished using as few as six individuals from each species (Blekhman et al. 2010).

Beyond its future applications, however, our panel has already yielded insights into the pluripotent state in chimpanzees and humans. Both at the transcriptional and epigenetic level, our iPSCs are remarkably homogeneous both within and between species, significantly more so than their precursors cells. This finding aligns with our current understanding of the reprogrammed pluripotent state as a complex, highly regulated state (Jaenisch and Young 2008), deviations from which are likely to result in loss of pluripotency and lineage commitment. We were able to identify nearly 5,000 genes that are differentially expressed between human and chimpanzee iPSCs, as well as nearly 2,000 differentially methylated regions between the two species. Many of the effect sizes associated with our identified DE genes and DMRs are small, and some are possibly of little biological consequence. Nevertheless, others are likely to be significant and functional in their *in vivo* developmental equivalents; indeed in many cases we find that these genes are in regions associated with developmental processes (supplementary tables 5 and 6).

PSCs have been used to study developmental pathways *in vitro* (for example, Paige et al. 2012; Rada-Iglesias et al. 2012; Wamstad et al. 2012; Xie et al. 2013). Although doubtlessly some optimization of existing differentiation protocols will be necessary, our panel of iPSCs makes it possible to carry out comparative developmental studies between humans and chimpanzees, and firmly test the hypothesis that changes in gene regulation and expression, especially during development, underlie phenotypic differences between closely related species, especially primates (Britten and Davidson 1971; Carroll 2005; Carroll 2008; Jacob 1977; King and Wilson 1975). In addition, we should be able to recreate and test the effect of inter-species regulatory changes in the correct cell type and species environment, enabling studies that cannot otherwise be performed (or are not feasible to perform) in humans and non-human primates. The use of panels of iPSCs including lines from both humans and non-human primates will thus allow us to gain unique insights into the genetic and regulatory basis for human-specific adaptations.

## Materials and Methods

### Isolation and culture of fibroblasts

All biopsies and animal care were conducted by the Yerkes Primate Research Center of Emory University under protocol 006-12, in full accordance with IACUC protocols. Skin punch biopsies (3 mm) were rinsed in DPBS containing Primocin (Invivogen) and penicillin/streptomycin (Pen/Strep, Corning) and manually dissected into 10-15 smaller pieces. The tissue was digested in 0.5% collagenase B (Roche) for 1-2 hours until cells were released from the extracellular matrix. Dissociated cells were pelleted by centrifugation at 250 x g, and the supernatant was spun a second time at 700 x g to pellet any cells that had not been completely released from the extracellular matrix. Cell pellets were resuspended in a 1:1 mixture of α-MEM and F12 (both from Life Technologies) supplemented with 10% FBS (JR Scientific), NEAA, GlutaMAX (both from Life Technologies), 1% Pen/Strep, 64 mg/L L-ascorbic acid 2-phosphate sesquimagnesium salt hydrate (Santa Cruz Biotech) and Primocin. Cells were plated in a single well of a 6-well plate coated with 4 µg/cm^2^ human fibronectin (BD Biosciences) and 2 µg/cm^2^ mouse laminin (Stemgent). Cultures were grown at 5% CO^2^/5% O^2^ until confluent and then split using 0.05% trypsin. For routine passaging cells were cultured at 5% CO^2^ and atmospheric oxygen in primate fibroblast media, which is the same as plating media but does not contain F12 base media.

### Generation of retrovirally-reprogrammed iPSC lines (a failed attempt)

We initially attempted to generate lines by retroviral transduction using through transfection with pMXs- vectors encoding the human *OCT3/4, SOX2, KLF4, L-MYC* and *NANOG* sequences (Addgene plasmids 17217, 17218, 17219, 26022 and 18115) as well as vectors encoding the MSCV- VSV.G envelope protein (Addgene plasmid 14888) and MSCV gag-pol (Addgene plasmid 14887). 15 µg of each vector was transfected into 293FT cells (Life Technologies) using Lipofectamine 2000 (Life Technologies) as directed by the manufacturer. We collected virus-containing supernatant from the 293FT cells 48 and 72 h after transfection and immediately used this viral media to transduce chimpanzee fibroblasts, alongside 10 ug/mL of polybrene (Sigma Aldrich H9268). To aid viral penetration, we centrifuged the cells at 1800 RPM for 45 minutes following each transduction. 24 h after the second transduction, we replaced the viral media with A-MEM + 10% FBS, NEAA and Glutamax. Transduced fibroblasts were allowed to recover for a further 2 days and then seeded on γ-irradiated, CF-1-derived mouse embryonic fibroblasts (MEF) at a density of 10,000 cells/cm^2^, and maintained in hESC media (DMEM/F12 supplemented with 20% KOSR, 0.1 mM NEAA, 2 mM GlutaMAX, 1% Pen/Strep, 0.1 mM BME and 25 ng/mL human bFGF) supplemented with 0.5 mM valproic acid (Stemgent) until day 14. We obtained iPSCs from 5 chimpanzees by using this protocol. Yet, when we performed quality control and pluripotency checks on these lines we found that the exogenous transfected genes are still expressed (in all lines). Pluripotency in these lines could not be maintained exclusively through endogenous expression. We discarded all 5 lines and proceeded with a different reprograming strategy as detailed below.

### Generation of episomally-reprogrammed iPSC lines

Fibroblasts were grown at 5% CO^2^/atmospheric O^2^ in primate fibroblast media until 70-80% confluence and released by trypsinisation for transfection. 1.5×10^6^ cells were transfected with 1.5 µg per episomal vector containing the following genes: *OCT3/4, SHp53, SOX2, KLF4, LIN28*, and *L-MYC* (Addgene plasmids 27077, 27078, 27080 and 27082; (Okita et al. 2011)). To boost the initial retention of vectors following transfection, 3 µg of *in vitro* transcribed ARCA capped/polyadenylated EBNA1 mRNA was cotransfected with the vectors (see below). Transfected cells were seeded at 15,000/cm^2^ on tissue culture plates precoated with 1 µg/cm^2^ vitronectin (Stemcell Technologies). Cells were grown in Essential 8 media (made in house as previously described in (Chen et al. 2011)) without *TGFβ1*, supplemented with 0.5 mM sodium butyrate (NaB, Stemgent)) and 100 nM hydrocortisone (Sigma Aldritch). Hydrocortisone was used between days 1-12, or until cell density exceeded >70% confluence. At day 12, cells were detached using TrypLE (Life Technologies) and replated at a density of 5,000 cells/cm^2^ on cell culture dishes precoated with 0.01 mg/cm^2^ (1:100) of hESC-grade Matrigel (BD Sciences) and grow in Essential 8 media without TGFβ1 or NaB. Colonies began to form at days 18-22 and were picked between days 24-30 onto dishes coated with γ-irradiated CF-1 derived MEF and subsequently grown in hESC media (as described above) supplemented with 100 ng/mL human bFGF (Milteny Biotech). Clones were routinely split using Rho-associated kinase (ROCK) inhibitor Y27632 (Tocris) at a concentration of 10 µM. Cells were migrated to 1:100 hESC Matrigel (BD Sciences) and maintained on Essential 8 media after a minimum of 15 passages on MEF. Feeder free cells were passaged using EDTA-based cell release solution as in (Chen et al. 2011).

### Generation of *EBNA1* mRNA

To generate a template for in vitro transcription, an EBNA1 template was designed using the wild type HHV4 *EBNA1* as a reference sequence (NCBI accession YP_401677.1). The reference sequence was modified by replacing the GA repeat region and domain B (amino acids 90-375) with a second, tandem, chromatin-binding domain (domain A, amino acids 27-89), similar to what was done by (Howden et al. 2006). The nuclear localization signal (amino acids 379-386) was removed and replaced with the sequence GRSS. Using the amino acid sequence as the starting template, the corresponding DNA sequence was generated by reverse translation and optimized for expression in human cell lines using Genscript’s OptimumGene codon algorithm. This sequence was synthesized by Genscript and provided in the pUC57 cloning vector; the EBNA1 coding sequence was subcloned into pcDNA3.1+ (Life Technologies) using the restriction enzymes BamHI and HindIII. Capped and poly(A) mRNA transcripts were generated using the mMESSAGE mMACHINE T7 ULTRA kit (Life Technologies) with 1 µg of BamHI linearized pcDNA3.1+EBNA1 as the template. The plasmids encoding the wild type and modified EBNA1 sequences have been deposited to Addgene as plasmid ID#s 59199 and 59198 for the wild type and modified sequences respectively.

### iPSC characterization

iPSC colonies were cultured on MEF for 4-6 days and fixed using PBS containing 4% PFA (Santa Cruz BioTech) for 15 minutes at room temperature. After rinsing with PBS, fixed cells were blocked and permeabilised for one hour in PBS containing 0.3% triton and 5% BSA. Primary antibodies: OCT3/4 (SC-5279), SOX2 (SC-17320), NANOG (SC-33759), SSEA-4 (SC-21704), and Tra-1-81 (SC-21706), all from Santa Cruz BioTech, were diluted 1:100 in blocking solution. Fixed cells were incubated with the primary antibody solution overnight on a rocker at 4 °C. After washing out the primary antibody solution, fixed cells were incubated with secondary antibodies (labeled with either Alexa-488 or Alexa-594, 1:400, Life Technologies) diluted in blocking for 1 hour on a rocker at room temperature. Nuclei were counterstained using 1 µg/mL Hoechst 33342 (ThermoFisher). All fluorescence imaging was conducted using an AMG EVOS FL (Life Technologies).

### Quantitative PCR for endogenous and exogenous gene expression

RNA was extracted using Qiagen RNA miniprep columns from cell pellets collected from fibroblasts, day 7 post transfection and feeder free (Matrigel and Essential 8) iPSC lines at passage 10 or higher for both the retroviral and episomal reprogrammings; 1 µg of total RNA was reverse transcribed using the Maxima first strand cDNA synthesis kit (Thermo Scientific). Quantitative PCR was performed using a 1:96 dilution of cDNA and SYBR Select master mix (Life Technologies) with both forward and reverse primers at a concentration of 0.2 µM.). Data was collected and analysed using the Viia7 (Life Technologies). Primer sequences are shown in supplementary table 2.

### Generation of embryoid bodies and immunofluorescence

Colonies growing on MEF were detached using Dispase/Collagenase IV (1 mg/ml each; both from Life Technologies) in DMEM/F12 and grown as a suspension culture on low adherent plates using hESC media without bFGF. After one week of suspension growth, cells were transferred to 12 or 24-well plates coated with 0.1% gelatin and grown in DMEM supplemented with 20% FBS, 0.1 mM nonessential amino acids, 2 mM GlutaMAX, 1% Pen/Strep and 64 µg/mL L-Ascorbic acid 2-phosphate sesquimagnesium salt hydrate. Embryoid bodies were grown for 1-2 weeks prior to fixation and immunofluorescence staining. Cultures were fixed and stained as described above using the following antibodies: AFP (1:200, SC-130302, Santa Cruz Biotech), FOXA2 (1:200, SC-6554, Santa Cruz Biotech), α-smooth muscle actin (1:1500, CBL171, Millipore) and MAP2 (1:200, sc-20172 and sc-74420, Santa Cruz Biotech).

### Integration analysis

To test for genomic integration and residual retention of episomal plasmids, each iPSC line was migrated to feeder free conditions and grown beyond passage 15 on hESC-qualified Matrigel (1:100 dilution, BD Biosciences) coated plates in Essential 8 media (Life Technologies). DNA was extracted from feeder free cultures using DNeasy Blood and Tissue Kits (Qiagen). PCR was performed using 100 ng of genomic DNA, an annealing temperature of 72°C and 25 cycles using primers designed to amplify a region common to all episomal vectors used (supplementary table 2). Genomic DNA (100 ng) isolated from day 7 cultures, and 1 pg of each episomal vector were used as positive controls. PCR products were run on a 1% agarose gel and visualised using ethidium bromide.

### Karyotyping

After 15 passages on MEF and hESC media, cells were migrated to 1:100 hESC Matrigel (BD Sciences) and maintained on Essential 8 media for upwards of 6 passages. Feeder-free adapted cells were sent to Cell Line Genetics Inc (Madison, WI) for karyotyping as described in (Meisner and Johnson 2008).

### Teratoma formation assays

In vivo developmental potential of the reprogrammed cell lines was examined. Monolayer iPSCs from three chimpanzee lines were grown on Matrigel (1:100) in E8 medium (Life Technologies) and collected by EDTA treatment (Life Technologies). Cells were counted and resuspended at a ratio of 1:1 cell volume to Matrigel and kept on ice until the injection. Six-week-old CB17.Cg-*Prkdc*^*scid*^*Lyst*^*bg-J*^/Crl immunodeficient male mice were obtained (Charles River Laboratories) and approximately one million iPSCs for each clone were injected into the testis-capsule. After five to eight weeks teratomas were isolated, weighed, measured, dissected, and fixed in 10% formalin. The specimens were embedded in paraffin, stained with hematoxylin and eosin, and analyzed by a histopathologist. All animal work was conducted under the approval of the Institutional Care and Use Committee of UCSD (Protocol# S09090).

In addition, live feeder free iPSC cultures maintained in Essential 8 media on Matrigel iPSCs from C4955D (passage 15+7) were provided to Applied Stem Cell Inc. (Menlo Park, CA) for teratoma analysis as previously described (Chen et al. 2012).

### Species of origin identity of teratoma samples

DNA was extracted from frozen teratoma tissue using DNeasy Blood and Tissue Kits (Qiagen). For teratomas derived from individual C4955, core sections were isolated from FFPE embedded teratomas tissue using a 3 mm dermal punch tool; DNA was extracted from core samples using a QIAamp DNA FFPE Tissue Kit (Qiagen). PCR was performed using universal mitochondrial primers ((Kocher et al. 1989) supplementary table 2) amplifying cytochrome b (*Cytb*, chimpanzee reference sequence NC_001643:bp 14233-14598) or the 12S ribosomal gene (12S, NC_001643:bp 484-915) with 250-500 ng of genomic DNA as the starting template. Two-step PCR was conducted with an annealing temperature of 50°C for 1 minute and an extension step at 72°C for 4 minutes for a total of 30 cycles. DNA was purified using a Wizard SV gel and PCR Clean-up kit (Promega); dye terminator cycle sequencing was conducted by the University of Chicago Comprehensive Cancer Center using 60 ng of purified PCR template and 4 μM of either the forward or reverse primer. Alignment to the chimpanzee, human (NC_012920) and mouse (NC_005089) reference sequences was accomplished using CLC Main Workbench 6.9 (Qiagen) and MUSCLE (Edgar 2004).

### Microarray genotyping and PluriTest

RNA from passage ≥15 iPSCs was extracted using the QIAGEN RNeasy kit according to the manufacturer’s instructions. Quality of the extracted RNA was assessed using an Agilent Bioanalyzer 2100 (RIN scores for all samples ranged from 9.9 to 10), and RNA was processed into biotinylated cRNA and hybridized to the HT12v4 array using standard Illumina reagents as directed by manufacturer. Arrays were scanned using an Illumina HiScan, and data processed using Illumina’s GenomeStudio software. Using these data, we carried out PluriTest as previously described (Müller et al. 2011). Additionally, we mapped all detected HT12v4 probe sequences (n = 46,297) to the chimpanzee (panTro3) genome using BWA 0.6.3 (Li and Durbin 2009). Probes that mapped to a single genomic location with no mismatches were retained (n = 21, 320, 46.2% of all probes) for the analysis that was restricted only to the chimpanzee lines.

When we considered data from human and chimpanzee iPSCs together, without excluding probes based on sequence matches to the chimpanzee genome, all chimpanzee lines in the panel had pluripotency scores slightly below the pluripotency threshold (supp figure 15, lighter points). However, low pluripotency scores could stem from differences in our ability to estimate gene expression levels in the chimpanzee compared to the human due to attenuated hybridization caused by sequence divergence (Gilad et al. 2005). Indeed, when we subset the array to retain only those detected probes that map to the chimpanzee genome with no ambiguity or mismatches, all chimpanzee lines have pluripotency scores greater than the pluripotency threshold value of 20 (supplementary figure 15, darker points).

### RNA sequencing and differential expression testing between iPSCs

50bp single-end RNA sequencing libraries were generated from RNA extracted from 7 chimpanzee and 7 human iPSC lines using the Illumina TruSeq kit as directed by the manufacturer (San Diego, CA), as well as from their precursor fibroblast or LCL cell lines. All iPSC samples were multiplexed and sequenced on four lanes of an Illumina HiSeq 2500; while the precursor cell lines were multiplexed and sequenced on six lanes of the same sequencer. We generated a minimum of 28,010,126 raw reads per sample (supplementary table 9), and confirmed the raw data were of high quality using FastQC (available online at http://www.bioinformatics.babraham.ac.uk/projects/fastqc/). We mapped raw reads to the chimpanzee (panTro3) or human (hg19) genome as appropriate using TopHat 2.0.8 (Trapnell et al. 2009), allowing for a maximum of 2 mismatches in each read. Due to the relatively poor annotation of the chimpanzee genome and to prevent biases in expression level estimates due to differences in mRNA transcript size and genetic divergence between the two species, we limited the analysis to reads that mapped to a list of orthologous metaexons across 30,030 ENSEMBL genes drawn from hg19 and panTro3, as in (Blekhman et al. 2010). Following mapping, gene level read counts were generated using *featureCounts* 1.4.4 as implemented in Subread (Liao et al. 2013). All sequencing data are available in the GEO database under series number GSE60996.

We normalized our data in two ways. In one instance, we retained only RNA-sequencing data from chimpanzee and human iPSCs, and retained 12,501 genes with at least 4 observations in one of the two species of log2 CPM > 1. CPM were then loess normalized by species within individuals with *voom* (Law et al. 2014). Finally, because the orthologous genes are not constrained to be the same length in both species, we computed RPKM for each gene before carrying out any inter-species comparisons. We then used the R/Bioconductor package *limma* 3.20.3 (Smyth 2004) to test for differential expression in our RNA-seq data. Finally, we tested for an enrichment of GO categories amongst DE genes using the R package *topGO* 2.16.0 (Alexa et al. 2006). These normalised values were used only to identify genes differentially expressed between iPSCs of the two species.

For the dataset containing RNA-sequencing data from iPSCs and their precursors, we again only retained 13,486 genes with at least 4 observations in one of the four groups (chimpanzee iPSCs, chimpanzee precursors, human iPSCs or human precursors) of log2 CPM > 1. Gene counts were then loess normalised within individuals by tissue, after correcting for the lack of independence within different tissues from the same individual, through the function corfit. As above, we then computed species-specific RPKM values, and used *limma* and *topGO* to test for differential expression and GO category enrichment, respectively. In this instance, we used a model design with 6 parameters (chimpanzee iPSC, human LCL-derived iPSC, human fibroblast-derived iPSC, chimpanzee fibroblast, human LCL and human fibroblast).

To confirm that our conclusions are robust with respect to the choice of normalization procedure, In both cases, we also tried a variety of other normalization schemes, including correcting for %GC content as in (Risso et al. 2011), none of which had a substantial effect on the final results (supplementary table 15). Finally, we built neighbor joining trees using Manhattan distances calculated from RPKM values at all 13,486 genes using the nj function in the R library *ape* (Paradis et al. 2004). All analyses were performed at a false discovery rate (Benjamini and Hochberg 1995) threshold of 1% unless otherwise noted, using R 3.1.0 (R Core Team 2013) and Bioconductor 2.14 (Gentleman et al. 2004).

### Methylation arrays

To analyze DNA methylation, we extracted DNA from all chimpanzee and human iPSC lines described above, as well as from the source fibroblast or lymphoblastoid cell lines. In all cases, 1000 ng of genomic DNA were bisulphite-converted and hybridized to the Infinium HumanMethylation450 BeadChip at the University of Chicago Functional Genomics facility as directed by the manufacturer. Since the probes on the array were designed using the human reference genome, we followed the approach described in (Hernando-Herraez et al. 2013) to compare humans and chimpanzees. We retained those probes that had either a perfect match to the chimpanzee reference genome, or had 1 or 2 mismatches in the first 45 bp but no mismatches in the 3’ 5 bp closest to the CpG site being assayed. We also removed all probes that contained human SNPs (MAF ≥0.05) and chimpanzee SNPs (MAF ≥0.15) within the last 5 bp of their binding site closest to the CpG being assayed. Within each individual, probes with a detection *P* > 0.01 were excluded. This resulted in the retention of 335,307 autosomal probes, and an additional 8,210 X chromosome probes, which we normalized and analyzed separately by sex. In all cases we performed a two-color channel signal adjustment, quantile normalization and β-value recalculation as implemented in the lumi package (Du et al. 2008). Because the HumanMethylation450 BeadChip contains two assay types which utilize different probe designs, we performed a BMIQ (beta mixture quantile method) normalization (Teschendorff et al. 2013) on the quantile-normalized autosomal data set. We did not perform this step on the X chromosome data, due to its methylation patterns. We built neighbor joining trees using Manhattan distances at all 335,307 probes using the nj function as above.

In order to identify differentially methylated probes we used an identical approach to that described above for the identification of DE genes. First, we identified probes that were differentially methylated between the iPSCs of both species using *limma* by using a reduced data set and model containing only data from the iPSCs themselves. Then, we fit a linear model to the data using limma with 6 parameters corresponding to the 6 tissue/species combinations in the data, classifying probes as differentially methylated at an FDR of 1%. As with the expression data, the reduced model has more power to identify DM probes between the two iPSC groups than the full model; however, there is great concordance between the two sets of results (supplementary figure 16). We excluded all probes with mean β inter-group differences < 0.1 in order to group DM probes into DMRs, which we define as 2 or more DM probes separated by < 1kb, with the additional requirement that the effect be in the same direction in all DM probes within the region. Finally, to examine the content of these DMRs, we used annotation files for the HumanMethylation450 Bead Chip provided by the manufacturer and discarded all DMRs associated with either multiple or no genes. We tested for enrichment of GO BP categories amongst the genes contained in the DMRs by using the R package topGO 2.16.0 (Alexa et al. 2006), using as a background set all genes in which it is theoretically possible to detect DMRs.

All methylation data is available in the GEO under series number GSE60996; A table with *P*-values for all hypothesis testing performed using the methylation data (by probe) is available on the Gilad lab website (http://giladlab.uchicago.edu/Data.html).

### Other indicators of genomic stability

Finally, we assessed two broad indicators of stability in our chimpanzee lines. All iPSC lines derived from female chimpanzees, and 3 of 4 lines derived from human females, show strong evidence for elevated expression of XIST relative to male lines (FDR-adjusted *P* = 0.0010; supplementary figure 17) and maintenance of X-chromosome inactivation during pluripotency. X-chromosome methylation patterns in females corroborate these observations, with the majority of probes mapping to the X-chromosome in our data being either hemimethylated (0.2 < β < 0.8) or hypermethylated (β ≥ 0.8) in females but not in males (supplementary figure 18). We also used a list of 168 imprinted probes from (Ma et al. 2014) to check for maintenance of genomic imprinting after reprogramming. We find that the majority of imprinted loci remain hemimethylated following reprogramming in both human and chimpanzee iPSC lines (supplementary figure 19). However, we identify two sets of probes that are consistently hypermethylated in pluripotent lines but were hemimethylated in their precursor cells. The first cluster contains 5 probes that are hypermethylated across both chimpanzee and human iPSCs; these probes are associated with the genes *KCNK9, ANKRD11* and *MKRN3*. The second cluster is comprised of 21 probes that are hypermethylated in all human iPSCs but only 2 chimpanzee iPSCs in our data, and is associated with the gene *PEG3-ZIM2*, which has been previously shown to be abnormally methylated in both hESCs and hIPSCs (Lund et al. 2012).

### Data Access

All novel RNA-sequencing and methylation data are available at the GEO under series number GSE60996. Additionally, a table with *P*-values for all hypothesis testing performed using the methylation data (by probe) is available on the Gilad lab website (http://giladlab.uchicago.edu/Data.html).

## Acknowledgements

We thank members of the Gilad and Marques-Bonet labs for helpful discussions; Julien Roux for help with differential gene expression analysis; Alexander Meissner for sharing methylation data from human iPSC and ESC lines and Fred Gage for providing a list of differentially expressed genes in retrovirally-reprogrammed chimpanzee iPSCs. This work was supported by NIH grant GM077959 to YG as well as by grants from the California Institute for Regenerative Medicine (CIRM); CL1-00502 and TR01250 to JFL, and ERC Starting Grant 260372 and MICINN (Spain) BFU2011-28549 to TM-B. IGR is supported by a Sir Henry Wellcome Postdoctoral Fellowship; NEB is supported by an NIH training grant (GM007197) and an NIH pre-doctoral award (F31 AG 044948); IHH is supported by FI Generalitat Catalunya.

## Supplementary figure legends

Supplementary figure 1: Karyotypes for the 6 chimpanzee iPSC lines not shown in main text figures, generated after >15 passages in culture. Passage number for each line represents passages on MEF feeders plus additional passages on Matrigel.

Supplementary figure 2: ICC staining of the 6 chimpanzee iPSC lines not shown in main text figures with antibodies for pluripotency markers as indicated. Scale bar: 200 μm.

Supplementary figure 3: ICC staining showing SSEA1 expression in chimpanzee iPSC culture plates, clearly distinct from NANOG expression.

Supplementary figure 4: Melt curves showing a lack of exogenous reprogramming gene expression in episomally reprogrammed chimpanzee iPSCs after > 10 passages.

Supplementary figure 5: ICC staining of differentiated embryoid bodies derived from the 6 chimpanzee iPSC lines not shown in main text figures, with antibodies indicating the three germ layers as indicated. Scale bar: 200 μm.

Supplementary figure 6: Histological staining of teratomas derived from three additional chimpanzee iPSC lines, showing generation of tissues from all three germ layers. Scale bar: 500 μm.

Supplementary figure 7: Sequencing traces from teratomas generated from chimpanzee iPSC lines for the mitochondrial genes 12S (C3649, C4955) and *cytb* (C8861, C40210). All traces show clear evidence of the presence of chimpanzee tissue in the teratoma.

Supplementary figure 8: Differentially expressed genes between iPSCs of chimpanzee and human origin.

Supplementary figure 9: Normalized mean expression values in chimpanzee and human iPSCS at the 73 differentially expressed genes associated with the 10 significantly overrepresented GO BP terms in the gene expression data. Red circles indicate a large group of ribosomal protein genes associated with most of these terms.

Supplementary figure 10: Boxplots of methylation beta values at 335,307 probes across all samples. Plots are colored by tissue type: light blue: chimpanzee iPSCs; dark blue: human iPSCs; light orange: chimpanzee fibroblasts; dark orange: human fibroblasts; turquoise: human LCLs.

Supplementary figure 11: Boxplots of methylation beta values across all samples, grouped by potency and genomic features. Boxes are colored by tissue type: light blue: chimpanzee iPSCs; light orange: chimpanzee fibroblasts.

Supplementary figure 12: Venn diagrams showing overlap in interspecies differences before and after reprogramming. a. Overlap in DE genes between chimpanzee and human fibroblasts, and chimpanzee and human fibroblast-derived iPSCs. b. Overlap in DM probes between chimpanzee and human fibroblasts, and chimpanzee and human fibroblast-derived iPSCs.

Supplementary figure 13: Venn diagram showing overlap of genes identified as DE between iPSCs of the two species when we normalize the iPSC data independently and alongside data from the precursors.

Supplementary figure 14: Exogenous gene expression in retrovirally reprogrammed chimpanzee iPSCs after various passages. All values are relative to expression in a day-7-post-transfection chimpanzee fibroblast.

Supplementary figure 15: The effects of probe sub-setting in PluriTest pluripotency score calculations. Lighter shades indicate pluripotency scores before the removal of probes not mapping to the chimpanzee genome, darker shades indicate pluripotency after probe removal. Purple circles denote chimpanzees; yellow squares, humans.

Supplementary figure 16: Venn diagram showing overlap of probes identified as DM between iPSCs of the two species under the full and reduced *limma* models.

Supplementary figure 17: Normalized expression values in 7 chimpanzee and human iPSCS at XIST. Circles denote chimpanzee iPSCs, squares indicate human iPSCS.

Supplementary figure 18: Quantile-normalized methylation beta values at 8,210 X-chromosome probes in 7 chimpanzee iPSCs and 7 human iPSCs. The colour bar beneath the dendrogram indicates sex of the individuals: purple: female; yellow: male. Sample names ending with _FB indicate fibroblast lines used to generate the corresponding iPSC line, samples ending with _LCL indicate LCL lines used to generate the corresponding iPSC line.

Supplementary figure 19: Normalized methylation beta values at 168 assayable probes known to be subject to parental imprinting effects, from (Ma et al. 2014). Sample names ending with _FB indicate fibroblast lines used to generate the corresponding iPSC line, samples ending with _LCL indicate LCL lines used to generate the corresponding iPSC line.

## Supplementary table headings

Supplementary table 1: Descriptive data for all chimpanzee cell lines used in this work.

Supplementary table 2: Origin and purpose of all primers used.

Supplementary table 3: Descriptive data for all human iPSC lines used.

Supplementary table 4: Normalized RPKM values and DE genes between chimpanzee and human iPSCs.

Supplementary table 5: Gene Ontology BP terms associated with genes DE between chimpanzee and human iPSCs.

Supplementary table 6: DMRs identified between chimpanzee and human iPSCs.

Supplementary table 7: Numbers of DM probes and DMRs between chimpanzee and human iPSCs identified under various mean β difference thresholds.

Supplementary table 8: Gene Ontology BP terms associated with genes within DMRs between chimpanzee and human iPSCS.

Supplementary table 9: RNA-sequencing reads generated and mapped for all samples in this work.

Supplementary table 10: Correlations between principal components and selected covariates in the expression data.

Supplementary table 11: Correlation between principal components and selected covariates in the methylation data.

Supplementary table 12: Normalized RPKM values and DE genes identified under the full *limma* DE testing framework.

Supplementary table 13: DMRs identified between chimpanzee iPSCs and their precursor fibroblasts.

Supplementary table 14: DMRs identified between human iPSCs and their precursor cells.

Supplementary table 15: Effects of different normalization schemes on the number of genes classified as DE in the full data set.

